# SHINE: Deterministic Many-to-Many clustering of Molecular Pathways

**DOI:** 10.1101/2025.02.07.636541

**Authors:** Lexin Chen, Jeremy M. G. Leung, Krisztina Zsigmond, Lillian T. Chong, Ramón Alain Miranda-Quintana

## Abstract

State-of-the-art molecular dynamics (MD) simulation methods can generate diverse ensembles of pathways for complex biological processes. Analyzing these pathways using statistical mechanics tools demands identifying key states that contribute to both the dynamic and equilibrium properties of the system. This task becomes especially challenging when analyzing multiple MD simulations simultaneously, a common scenario in enhanced sampling techniques like the weighted ensemble strategy. Here, we present a new module of the MDANCE package designed to streamline the analysis of pathway ensembles. This module integrates n-ary similarity, cheminformatics-inspired tools, and hierarchical clustering to improve analysis efficiency. We present the theoretical foundation behind this approach, termed Sampling Hierarchical Intrinsic N-ary Ensembles (SHINE), and demonstrate its application to simulations of alanine dipeptide and adenylate kinase.

## 1 INTRODUCTION

As we enter the exascale era of supercomputing, equipped with advanced simulation methods, generating atomistic pathways for functionally interesting processes has become increasingly practical in the sciences. While these pathways offer rich mechanistic insights, analyzing them can be a major challenge given their diversity and variable lengths not to mention the massive amounts of simulation data generated— ranging from tens to hundreds of terabytes. Consequently, a key objective for simulators is to develop software tools that facilitate mechanistic pathway analysis through rapid and reliable clustering into distinct, informative routes.

To this end, several path clustering methods have been developed. A common feature of these methods is the pairwise comparison of conformations (frames) among sampled pathways. Such comparisons have been performed using Hausdorff or Frechet geometric distances (pathway similarity analysis, PSA, and pathway histogram analysis, PHAT^1^), deep-learning models (latent-space clustering, LSP^2^), and linguistics-assisted clustering (Linguistics Pathway Analysis of Trajectories with Hierarchical clustering, LPATH^3^).

An alternate approach to clustering pathways involves considering all frames (or at least, a representative subset) of sampled pathways. Compared to clustering methods relying on pairwise frame comparisons, this approach incurs only modest additional computational costs when using recently introduced *n*-ary similarity functions. These functions have applications spanning drug-design, chemical space exploration, and imaging mass spectrometry. In molecular dynamics (MD) studies, one of the simplest and most widely used *n*-ary similarity measures is the mean squared deviation (MSD). A key advantage of MSD is that it quantifies the similarity between *N* frames with an O(*N*) memory and time cost.^4^

In this work, we present Sampling Hierarchical Intrinsic *N*-ary Ensembles (SHINE), a method that leverages the cost-effective MSD metric to quantify pathway distinctions. SHINE is inspired by LPATH, incorporating a similarity matrix that captures all pairwise pathway comparisons. It then applies Hierarchical Agglomerative Clustering with Ward linkage to process trajectory data. To further reduce computational costs, we integrate cheminformatic-inspired sampling algorithms^5^ to generate a distance matrix between trajectories, which can then be clustered using any hierarchical agglomerative clustering method. We demonstrate the effectiveness of SHINE in simulations of conformational transitions in two benchmark systems: alanine dipeptide and adenylate kinase.^3,6^ Our method is implemented as the SHINE module in the MDANCE post-processing package (https://github.com/mqcomplab/MDANCE).

## 2 THEORY

### A. *N*-ary Indices

A key feature of SHINE is its ability to calculate the similarity (or dissimilarity) between elements of a set in linear time and memory. Traditionally, for a set of *N* elements, calculating pairwise distances and averaging them requires O(*N*^2^) effort, as it involves comparing all possible pairs with 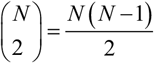 pairs. However, recent advances have introduced more efficient O(*N*) solutions to this problem^4,7–12^. In drug-like molecule studies, these advances led to the development of instant similarity indices, iSIM^13^, (or the extended indices, eSIM^5,14^). Meanwhile, in biomolecular analysis, the Mean Squared Deviation (MSD) has been widely used due to its close relationship with the commonly employed root mean squared deviation (RMSD). Despite their apparent differences, both iSIM and MSD share the same initial steps. First, the objects to be compared must be encoded as fixed-length vectors *F*. The *i*th object, *F*^(*i*)^, is represented by a one-dimensional array 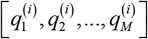 where 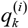 are the components used to represent the molecules or conformations (in the present study, as Cartesian or internal coordinates). Next, these object representations are arranged into an *N* by *M* matrix, where *N* is the number of objects, *M* is the number of coordinates. The key insight is that quantifying the relationships between these objects requires only two vectors: *S*, the column-wise sums of the representation matrix; and *R*, containing the column-wise sums of the squares of the elements of the representation matrix. Mathematically:

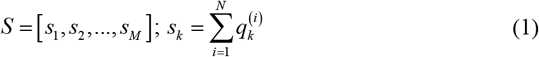

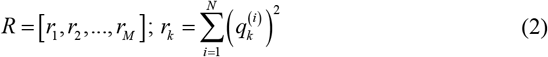

These vectors can be computed in O(*N*) time. They serve as the only necessary components for calculating either iSIM or MSD:

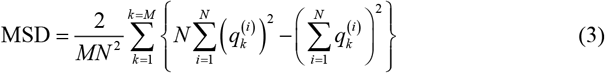

Large MSD values indicate more dispersed (less similar) sets, while smaller MSD values indicate similarity among conformations. SHINE supports the use of both MSD (default) and iSIM for similarity calculations in trajectory analysis.

### B. Frame Selection

A key challenge in the meaningful clustering of pathways is the selection of frames (conformations) that prevents “loops”— where pathways involving the same initial and final conformational states—are not mistakenly clustered into distinct pathway classes. To address this, LPATH employs a two-step procedure: (i) discretizing pathways as text strings by assigning frames (sampled at fixed time intervals) to distinct conformational states, and (ii) removing repeating frame patterns up to a specified limit using the “condense” option. SHINE tackles this challenge using two alternative strategies that do not require discretization: diversity selection and representative selection.

*Diversity selection*. The diversity picking problem is based on a simple premise: pick a subset that is maximally dissimilar. In SHINE, we use this algorithm: a) If no conformations have been selected, pick a representative from the trajectory and add it to the *Selected* set. b) The next molecule to be added will be the one that maximizes the MSD of the *Selected* set. This recipe guarantees to pick conformations from different sectors of the PES.

*Quota selection*. Most diversity-oriented sampling methods tend to prefer picking points from the periphery of the set, so it seems convenient to have a sampling method that picks points in a more uniform way. We will use our recently proposed quota sampling, which is based on the concept of complementary similarity. While the MSD of a set is a global measure, the complementary similarity (cMSD) is a local tool. The cMSD of a conformation is just the MSD of the set, after removing that conformation. This provides a convenient (O(*N*)) ranking of all the frames: bigger (lower) cMSD values correspond to “central” (“outlier”) points. We can use this to select points uniformly: a) Calculate the cMSD for all frames in the simulation, b) Order the frames according to their cMSD, c) Assign frames to bins, such that the difference between the highest and lowest cMSD values in each bin is constant, d) Pick a frame per bin. This guarantees that the space of frames is explored without biasing any particular region.

SHINE also provides an option for a “minimum trajectory size”. That is, the user could indicate that if a trajectory has no more than a desired number of frames (by default, 50 frames), no sampling will be performed for that trajectory.

### C. Pathway Similarity and Clustering

The final two steps of SHINE involve quantifying the relationships between trajectories and then using these relationships as input for a clustering algorithm. For clustering, we follow the approach used in PSA and LPATH by employing hierarchical agglomerative clustering (HAC). Specifically, we utilize the scipy.cluster functionality,^15^ which offers flexibility in handling the HAC.^16^ As recommended in PSA and LPATH, we favor the Ward linkage criterion in the agglomerative steps, since it has been shown to provide more robust results.^17^ The optimum number of clusters (distinct pathways) is determined by identifying the largest change in inter-cluster distance.

This raises a key question: how should trajectory similarity be measured? As noted above, we aim to use methods that are as direct as possible, requiring minimum data pre-processing. Additionally, we prioritize computational efficiency.

A common assumption in trajectory comparisons is that the triangle inequality must hold. However, while this is essential while comparing individual points in a metric space, here we are not comparing single points (conformations), but rather sets of points (trajectories). Since we do not know whether these sets exist within a well-defined metric space, we opt to relax the triangle inequality constraint. As a result, our trajectory comparisons do not adhere strict metric properties; instead, we refer to them as dissimilarity functions rather than true distance metrics.

We impose only two requirements on these dissimilarity functions:

1. **Symmetry:** The dissimilarity between trajectory A and trajectory B must be the same in either direction: *d* ( *A, B*) = *d* ( *B, A*).
2. **Relative ranking:** The results must provide a ranking of how close or far apart trajectories are. In other words, if *d* ( *A, B*) < *d* ( *A, C*), we infer that trajectory A is more similar to trajectory B than to trajectory C.

Remarkably, we do not require the dissimilarity values to be strictly positive. Since we only care about the relative ranking of these values, these two cases: a) *d* ( *A, B*) = 1, *d* ( *A, C*) = 2, b) *d* ( *A, B*) = −3, *d* ( *A, C*) = 7 carry the same information. This flexibility allows us to explore new inter-trajectory dissimilarities while also improving computational efficiency. For instance, given two trajectories A and B, with *N*_*A*_, *N*_*B*_ frames, respectively, computing the Hausdorff metric between them scales as *O* ( *N*_*A*_*N*_*B*_), which quickly becomes unmanageable. However, this complexity is unavoidable if one insists on preserving the triangle inequality.

In SHINE we provide the option to use the Hausdorff metric (calculated from the scipy.spatial.distance module) for users who require triangle inequality-abiding results. However, we also provide a broader range of more computationally efficient alternatives:

1-Intra dissimilarity:

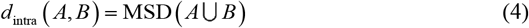

2-Inter dissimilarity:

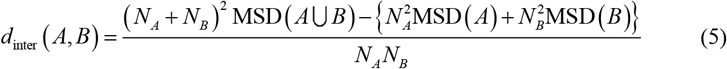

3-Semi-sum:

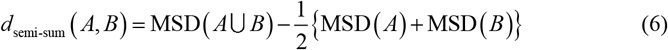

4-Min:

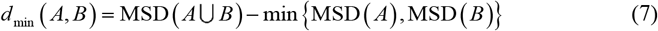

5-Max:

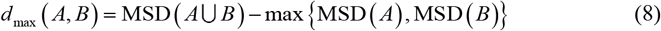

All these functions scale as *O* ( *N*_*A*_ + *N*_*B*_). *intra* is the simplest, since it just quantifies the separation between A and B by the MSD of the union of the two trajectories. *Inter* resembles the average linkage criterion, since it only takes into account the MSD values of points in A and B. This is similar to *semi-sum*, which weighs the relative importance of MSD(A) and MSD(B) in a different way. Finally, *min* and *max* are inspired by the HAC stopping criterion proposed in Eq. (2).^17^

The SHINE workflow is summarized in Fig. 1.

**Figure 1:**
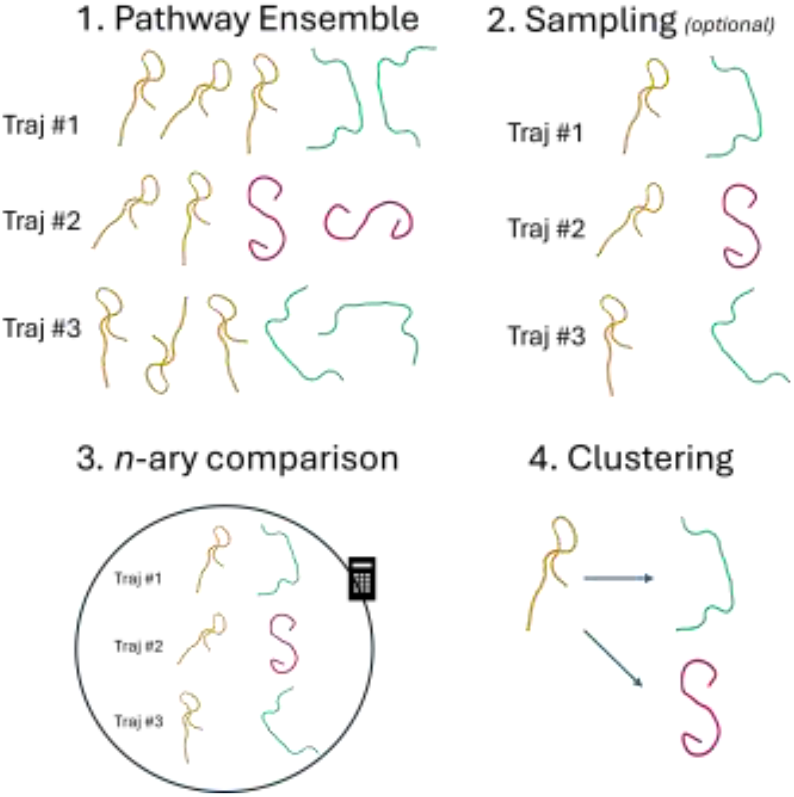
Schematic representation of the SHINE workflow, from the generation of the different simulations, to the (optional) sampling, the formation of the pairwise matrix of *n*-ary comparisons between the sampled conformations, and the final hierarchical clustering.

**Figure 2:**
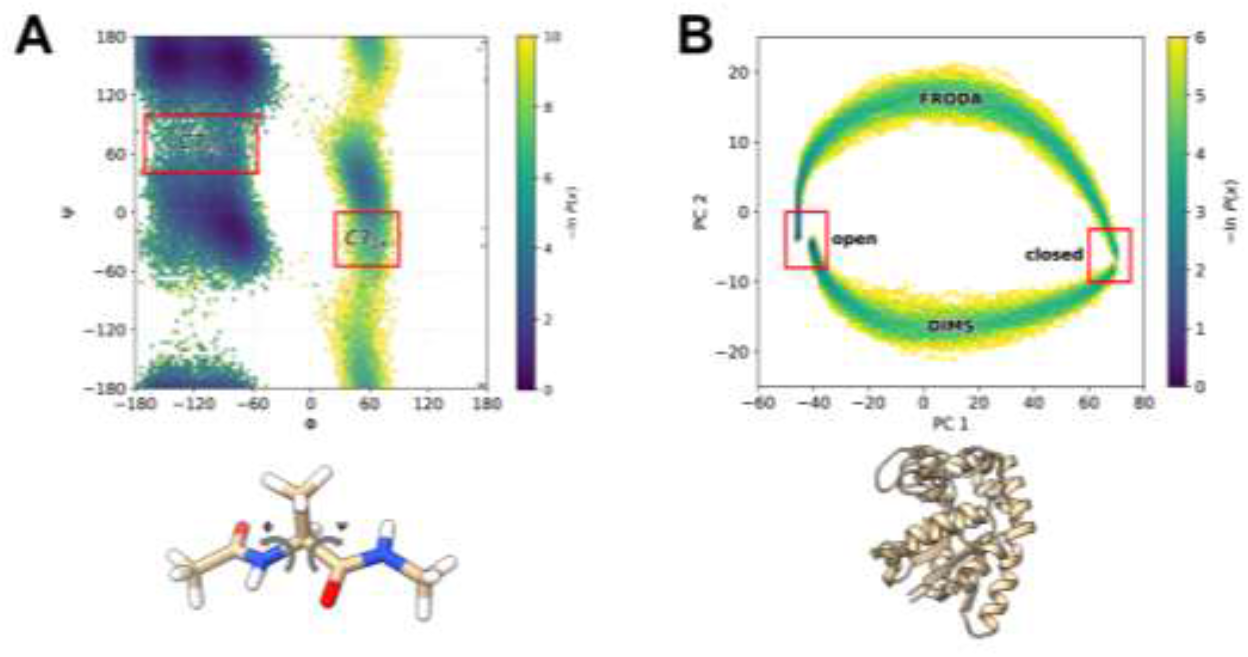
Benchmark systems and key conformational states, as highlighted in probability distributions from molecular simulations. **A**. Probability distribution of alanine dipeptide as a function of φ and Ψ backbone torsional angles. The slowest conformational transition is from the C7eq to C7ax conformational states. **B**. Probability distribution of adenylate kinase as a function of the first two principal components from principal component analysis (PCA) on xyz coordinates of all C_ª_ atom after alignment to the C_ª_ CORE domain^1,6^. Trajectories between the closed and open conformations of adenylate kinase were studied. The top and bottom paths in the 2D probability distribution contain structures sampled by FRODA^18^ and DIMS,^19^ respectively.

## 3. SIMULATION DETAILS

To demonstrate the effectiveness of SHINE, we applied the method to simulations of two benchmark systems: alanine dipeptide and adenylate kinase. Full simulation details for each system are provided below.

### Alanine dipeptide

An ensemble of 80 successful pathways for the slowest conformational transition of alanine dipeptide (C7eq to C7ax transition) were extracted from five independent WE simulations. WE simulations were run using the WESTPA 2.0 software package^20^ C with a resampling time interval **τ** of 100 ps and a two-dimensional progress coordinate consisting of the φ and Ψ backbone torsional angles. Fixed bins were positioned only along the Ψ dimension of the progress coordinate at 20° intervals between 0° and 360° with a target of 8 trajectories/bin. Dynamics in the WE simulations were propagated using the Amber 22 software package with the Amber ff14SBonlysc force field^21,22^ and generalized Born implicit solvent (igb=1).^23^ The hydrogen mass repartitioning scheme was applied to enable a 4-fs time step. Our set of five WE simulations yielded an aggregate simulation time of 14.6 µs with coordinates saved every 4 ps for analysis. Each WE simulation was completed in 21 h using 8 CPU cores of a 3.5 GHz Intel Xeon CPU in parallel.

### Adenylate kinase

An ensemble of 400 successful pathways involving the close-to-open transition of adenylate kinase from *Escherichia coli* was provided by Beckstein and co-workers.^1,6^ Half of these pathways were generated using the CHARMM c36b2 dynamics engine^24^ and dynamic importance sampling molecular dynamics (DIMS),^19^ a perturbation MD method using biased random walks in configurational space to sample towards the target structure.^6^ The remaining 200 pathways were generated in the absence of solvent using the Framework Rigidity Optimized Dynamics Algorithm (FRODA), a geometric targeting technique used to optimize stereochemical constraints and hydrophobic contacts. The DIMS simulations employed the CHARMM 22/CMAP protein force field, ACS/ACE2 implicit solvent model, and Langevin dynamics at 300K.

## 4. RESULTS

The SHINE analysis of the alanine dipeptide and the adenylate kinase in implicit solvent, gave consistent results. In both cases, the semi-sum and intra dissimilarity metrics are robust, with consistently accurate results over diverse calculation conditions (selection method and % of selected frames). Surprisingly, the min dissimilarity also performs quite well across various choices of the calculation parameters. On the other hand, the Hausdorff, inter, and max dissimilarities often lead to very disappointing results.

The LPATH analysis of the 80 successful weighted ensemble pathways for the C7eq to C7ax conformational transition of alanine dipeptide (labeled with indices [0-79]) resulted in two distinct classes (clusters) of pathways, with simulations [25-27, 33-47, 51, 68] in the least populated class. First, we studied the quota method. With this option, the Hausdorff, inter, and max dissimilarities failed to reproduce this cluster distribution. However, the semi-sum, min, and intra dissimilarities performed much better (see Fig. 3). Notice that (essentially independently of the fraction of sampling), semi-sum and intra quickly gather [33-47] in one cluster. Min, on the other hand, favors the [33-42, 47, 53] grouping. In almost all these cases, the optimum number of clusters is 2, with the exception of the semi-sum similarity without any sampling, that identifies 3 optimal clusters. What is interesting about this case is that it contains the [33-47] group and a [0, 7, 21, 25-27, 51, 68] cluster that, if combined, result in the smallest LPATH cluster.

**Figure 3:**
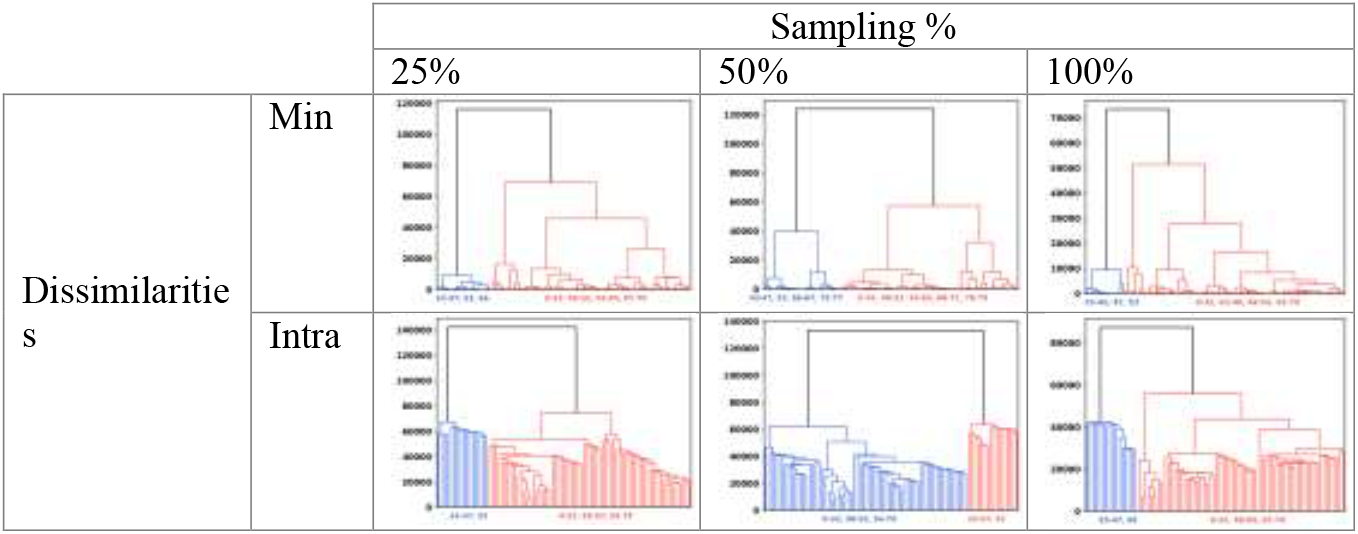
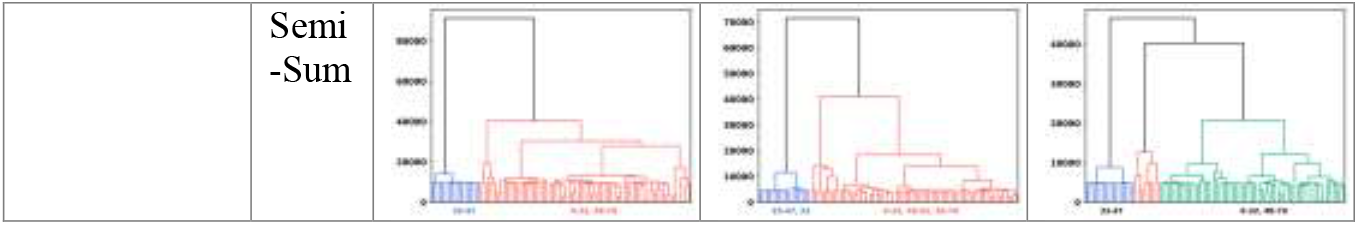
Dendrograms for the min, intra, and semi-sum dissimilarities at 25%, 50%, and 100% quota sampling of the ala-dipeptide trajectories.

The diversity selection of the alanine dipeptide simulations more or less follows the same trends of the quota selection (Fig. S1, SI). Once again, inter, max, and Hausdorff fail to capture the intrinsic structure of the ensemble of pathways. But now, the min dissimilarity is more well-behaved, recognizing the [33-47] cluster for a wide array of sampling fractions. What is remarkable about this approach is that the intra and semi-sum dissimilarities not only recognize the [33-47] group as well, but that they do so for all sampling fractions, from 5% to 100%.

Up to this point, we have been concerned with the problem of clustering pathways. However, inspired by the traditional clustering analysis of MD simulations, we can also ask the question of how to select representative trajectories from a given cluster. This typically involves selecting the *centroid*, or the average of coordinates for all structures in the cluster, as the representative structure of the cluster. However, just taking this average can lead to unphysical structures. Moreover, in the analysis of trajectories, it is not even clear how one should take this average, given that different simulations are not required to have the same number of frames. The solution to both these problems is to focus on the *medoid* of the set, as opposed to the centroid, with the key difference being that the medoid is required to be an element of the set. This is the approach that we take to analyze the SHINE clusters, with the added advantage that the most time-consuming step to calculate the medoid, generating the pairwise dissimilarity matrix between trajectories, was already performed. Then (using Python lingo), one just needs to take slices from the dissimilarity matrix using the indices of each cluster, and calculate the vector with the sums of the columns (or rows) of these sub-arrays, where the resulting minimum value of each set of vectors will correspond to the medoid of each cluster. In Fig. 4, we show this analysis for the alanine dipeptide. Here, we only considered the semi-sum, and we calculated the medoids of the top two clusters at 96 different values of sampling (from 5 to 100%), using the quota and diversity methods. It is reassuring to see that these different combinations produce consistent results, with pathways 44-45 and 67 being consistently selected as the most representative.

**Figure 4:**
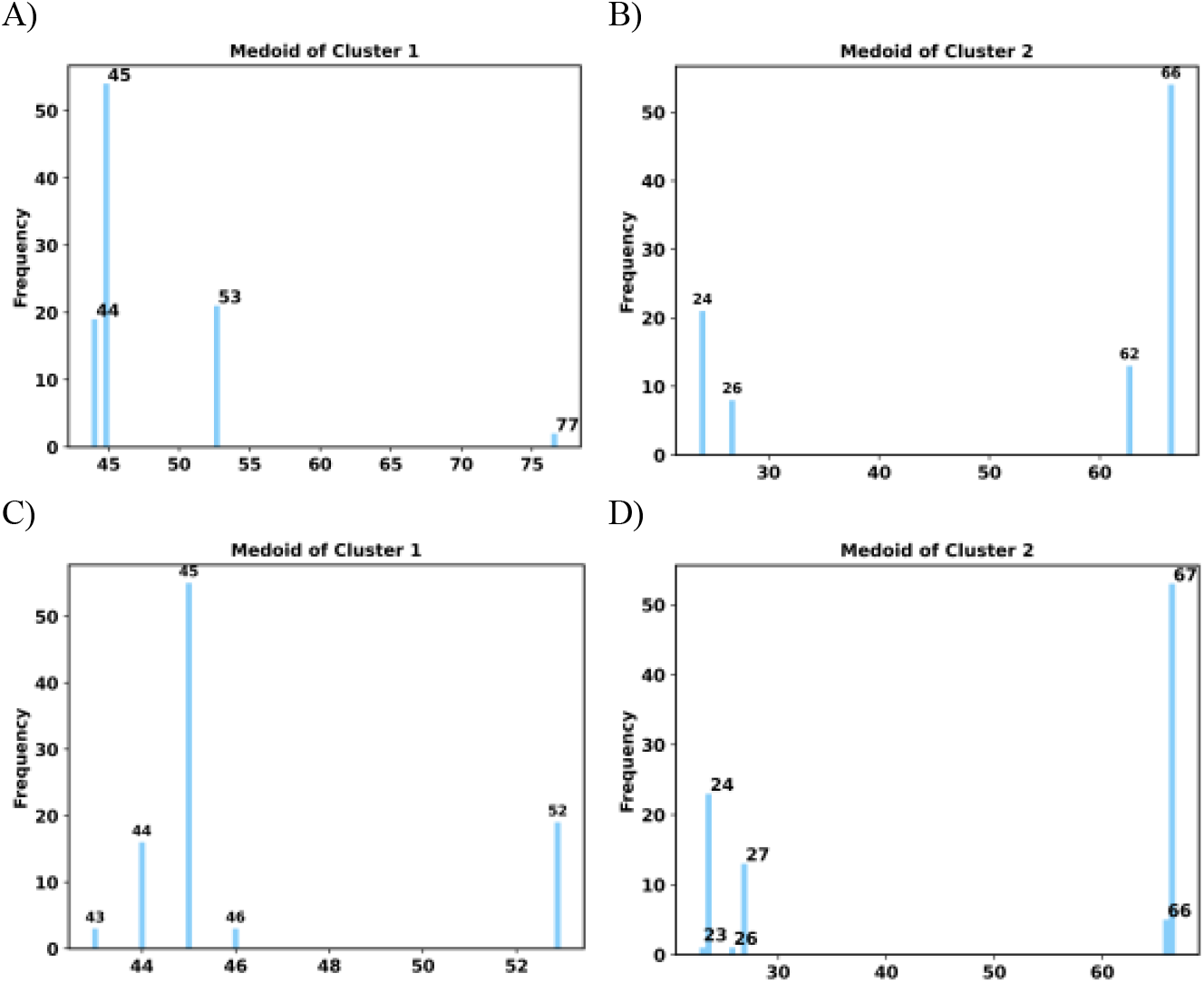
Most representative pathways in the two-clusters cases for the alanine dipeptide simulations after diversity (A, B) and quota (C, D) sampling. Semi-sum dissimilarity.

**Figure 5:**
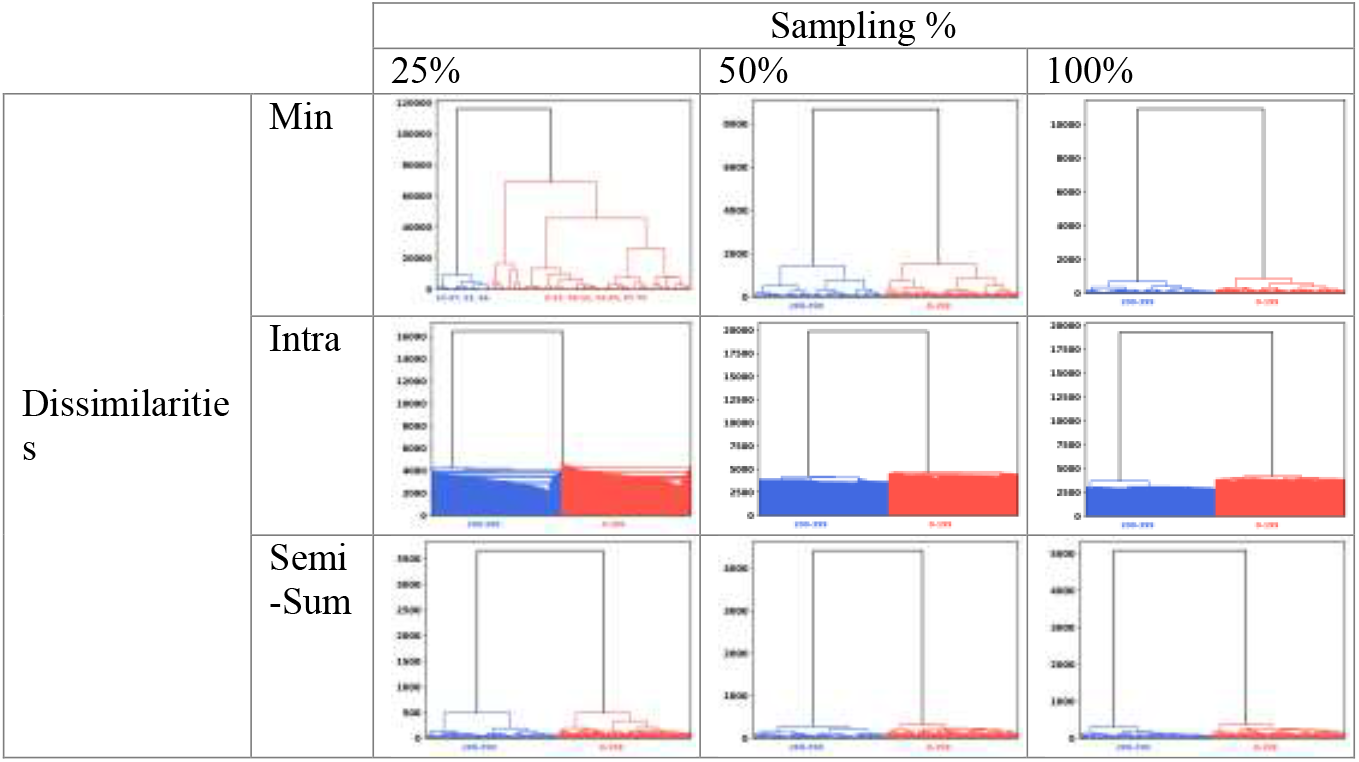
Dendrograms for the min, intra, and semi-sum dissimilarities at 25%, 50%, and 100% quota sampling of the adenylate kinase trajectories.

To showcase the generality and broad applicability of SHINE, we also considered a more complicated system: the close-to-open conformational transition in adenylate kinase (see Figs. 2). Here, we have 400 pathways for the conformational transition with indices [0-199] corresponding to FRODA and [200-399] to DIMS. As noted by Seyler et al.^1^, the Hausdorff metric succeeds in classifying these two types of simulations. However, the inter and max dissimilarities fail in recognizing this pattern, independently of the way of sampling the trajectories. On the other hand, min-, intra-, and semi-sums provide robust results in accordance with the PSA study. In this application, the diversity sampling seems to have a decisive advantage. For instance, the min dissimilarity only agrees with PSA if one samples more than 40% of the frames using quota sampling. This is improved upon by the intra method, which provides the correct answer if one samples > 20% of frames (again, with the quota method). Only the semi-sum dissimilarity with quota sampling shows perfect agreement with PSA, independently of the fraction of frames. However, if one uses the diversity sampling, all of min, intra, and semi-sum recover the PSA structure at any degree of sampling.

Finally, the medoid analysis of the adenylate kinase trajectories shows some differences between the different sampling methods (Fig. S3). Here, interestingly, the diversity sampling for “cluster 1” (indices in the [200-399] range) gives 6 potential candidates with counts close to or above 10, with 2 potential candidates for “cluster 2” ([0-199]) also with more than 10 counts. On the other hand, the quota sampling is more consistent at picking medoids: there are only 3 candidates for “cluster 1” and 1 for “cluster 2” with more than 10 counts.

## 5. CONCLUSIONS

SHINE offers an effective alternative to pathway clustering analysis based on *n*-ary comparisons. The many-to-many nature of *n*-ary indices makes them a natural choice for this task, enabling multiple possible definitions of pathway “separation”. SHINE achieves this without requiring extensive pre-processing and at a lower computational cost than previous approaches, particularly those relying on the Fréchet distance. Notably, our results indicate that enforcing the triangle inequality in the space of full simulations provides little benefit. While this may initially seem counterintuitive, it is important to recognize that although triplets of points satisfy this axiom under any metric, it is not a necessary condition for comparing multiple points simultaneously. Similar to other pathway clustering methods, a feature of SHINE is its ability to identify the most representative pathway within each pathway class (cluster). This capability provides valuable insight into the dominant mechanisms underlying diverse biomolecular transformations while simplifying the structural analysis of these paths. We demonstrate the effectiveness of SHINE through its application to simulated ensembles of pathways for conformational transitions in alanine dipeptide and adenylate kinase. Future studies will extend SHINE’s application to more complex biomolecular processes.

## Acknowledgements

RAMQ, LC, and KZ thank support from the National Institute of General Medical Sciences and the National Institutes of Health under award number R35GM150620. This work was also supported by NIH grant R01 GM1151805 to LTC and a MolSSI Software Fellowship to JMGL.

## Note

LTC serves in the Scientific Advisory Board of OpenEye Scientific Software.

