## Supplementary Information for "SHINE: Deterministic Many-to-Many clustering of Molecular Pathways"

### Many-to-many clustering of molecular pathways through Sampling Hierarchical Intrinsic *N*-ary Ensembles (SHINE)


| | | Sampling % | | |
|---|---|---|---|---|
| | | 25% | 50% | 100% |
| Dissimilarities | Min | 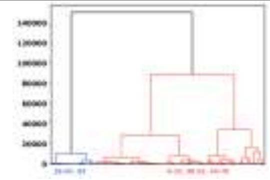 | 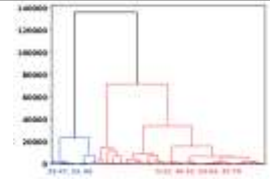 | 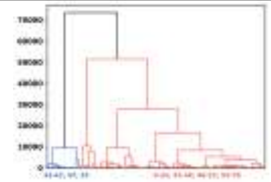 |
| | Intra | 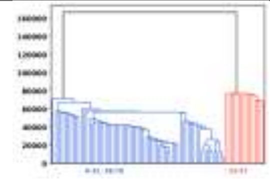 | 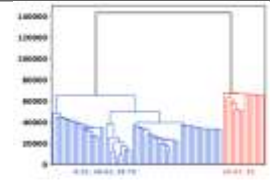 | 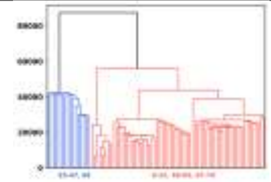 |
| | Semi-Sum | 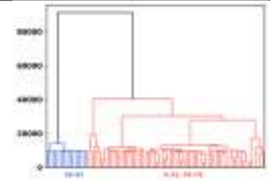 | 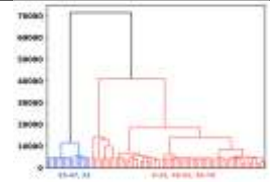 | 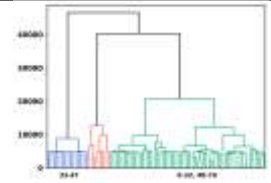 |

**Figure S1:** Dendrograms for the min, intra, and semi-sum dissimilarities at 25%, 50%, and 100% diversity sampling of the ala-dipeptide trajectories.

| | | Sampling % | | |
|---|---|---|---|---|
| | | 25% | 50% | 100% |
| Dissimilarities | Min | 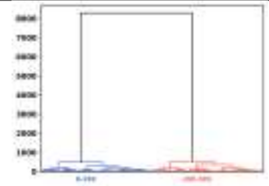 | 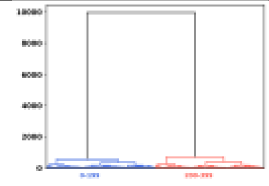 | 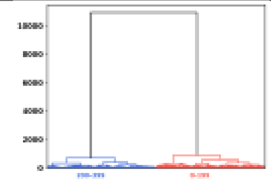 |
| | Intra | 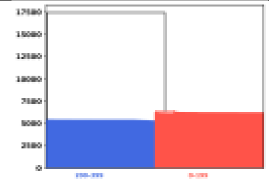 | 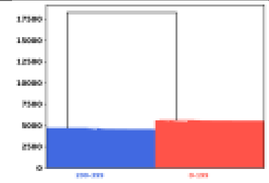 | 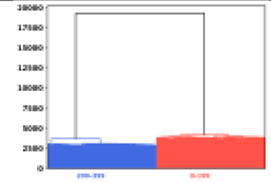 |
| | Semi-Sum | 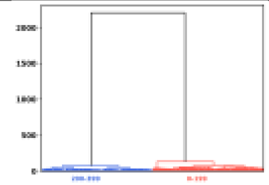 | 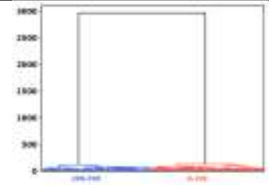 | 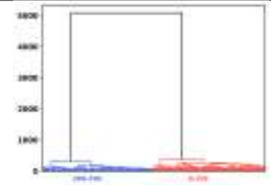 |

**Figure S2:** Dendrograms for the min, intra, and semi-sum dissimilarities at 25%, 50%, and 100% diversity sampling of the adenylate kinase trajectories.

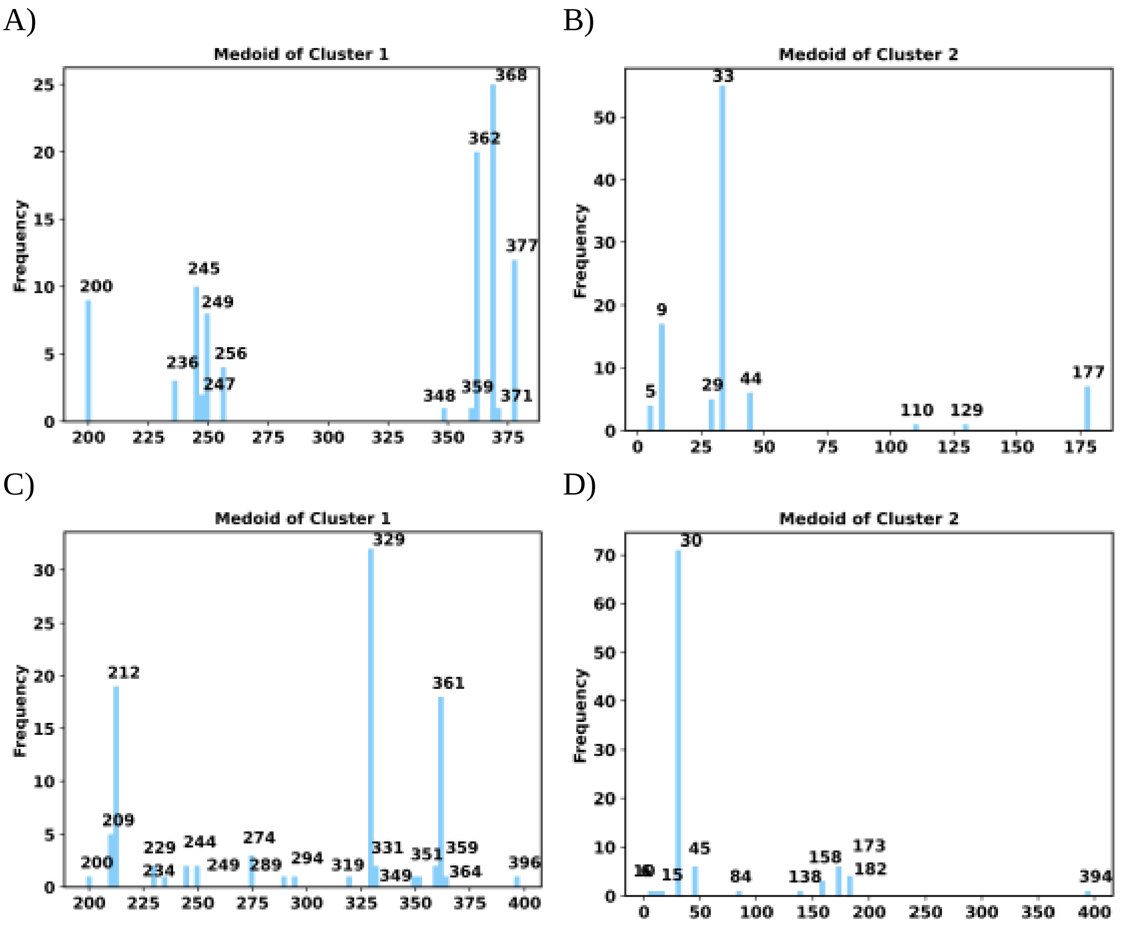


**Figure S3:** Most representative pathways in the two-clusters cases for the adenylate kinase simulations after diversity (A, B) and quota (C, D) sampling. Semi-sum dissimilarity.
